# Phylogenetic analyses suggest independent origins for trichromatic color vision in apes and Old World monkeys

**DOI:** 10.1101/2020.03.23.003855

**Authors:** Jessica Toloza-Villalobos, Juan C. Opazo

**Affiliations:** Instituto de Ciencias Ambientales y Evolutivas, Facultad de Ciencias, Universidad Austral de Chile, Valdivia, Chile; Millennium Nucleus of Ion Channels-associated Diseases (MiNICAD)

**Keywords:** trichromatic color vision, opsin gene, gene duplication, primate color vision

## Abstract

In catarrhine primates, trichromatic color vision is associated with the presence of three opsin genes that absorb light at three different wavelengths. The OPN1LW and OPN1MW genes are found on the X chromosome. Their proximity and similarity suggest that they originated from a duplication event in the catarrhine ancestor. In this study we use the primate genomes available in public databases to study the duplicative history of the OPN1LW and OPN1MW genes and characterize their spectral sensitivity. Our results reveal a phylogenetic tree that shows a clade containing all X-linked opsin paralogs found in Old World monkeys to be related to a clade containing all X-linked opsin paralogs identified in apes, suggesting that routine trichromacy originated independently in apes and Old World monkeys. Also, we found spectral variability in the X-linked opsin gene of primates. Our study presents a new perspective for the origin of trichromatic color vision in apes and Old World monkeys, not reported so far.

## Introduction

Trichromatic color vision is associated with the presence of three genes, called opsins. These genes encode for proteins expressed in the cone photoreceptor cells, which absorb light at three different wavelengths^1,2,3,4,5,6^. The short wavelength sensitive gene (OPN1SW) is located on an autosome, while the medium wavelength sensitive gene (OPN1MW) and long wavelength sensitive gene (OPN1LW) are separated by only 24 kb on the X chromosome^7,8^. The proximity and similarity of the OPN1LW and OPN1MW genes suggests a relatively recent duplication event from a single ancestral gene^8,9^.

Primates are unique among mammals in having trichromatic color vision, a phenotype that arose through several evolutionary pathways^7,10^. Catarrhine primates, the group that includes apes and Old World monkeys, possess routine trichromacy, a condition characterized by the presence of three opsin genes that absorb light at three different wavelengths^11,12^. From an evolutionary perspective, and given that duplicated X-linked opsin genes are only present in the genome of catarrhine primates, it has been proposed that the duplication event that gave rise to routine trichromacy occurred after the divergence from New World monkeys^3^. Although New World monkeys possess only one opsin gene on the X-chromosome, they have different types of vision^2,13^. Nocturnal monkeys, *Aotus trivirgatus*, possess monochromatic vision as they have only one functional opsin gene, located on the X chromosome^14,15,16^. Similar to catarrhine primates, howler monkeys of the genus *Alouatta* possess routine trichromacy. However, their OPN1LW and OPN1MW opsin genes are the product of an independent duplication event^13,17,18^. In the other members of the group, males have dichromatic color vision while some females have trichromatic color vision due to a polymorphism which produces an allele sensitive to medium wavelengths of the visual spectrum^6,12^. Thus, homozygous females have dichromatic color vision, while heterozygous females possess trichromatic color vision. Genetic assessment in tarsiers have revealed that different species possess different allelic forms^19^. In the case of the Bornean tarsier (*Tarsius bancanus*), seven medium wavelength sensitive alleles (M) are described, whereas in the Philippine tarsier (*Carlito syrichta*), 25 long wavelength sensitive alleles (L)^19^ are described (Tan and Li, 1999). Thus, tarsiers have dichromatic color vision. However, there are differences in color perception depending on the visual phenotype (red or green) of the X-linked opsin allele of the species^20^. Among strepsirrhines, monochromacy is described in nocturnal species, while in other species with a functional OPN1SW gene, allelic variation has been shown to give rise to allelic trichromacy^19^.

In this study we use the genomes available in public databases to study the duplicative history of the X-linked opsin genes in primate evolution and characterize their spectral sensitivity. Our phylogenetic tree showed a monophyletic group containing all X-linked opsin paralogs found in Old World monkeys to be linked to a monophyletic group containing all X-linked opsin paralogs identified in apes, suggesting that routine trichromacy originated independently in apes and Old World monkeys. We found variation in the spectral sensitivity of the X-linked opsin genes with seven maximum wavelengths of absorption. Among apes, we found two M alleles and three L alleles. New World monkeys, tarsiers and strepsirrhines, groups characterized by having only one X-linked opsin gene, also possess allelic variation. In Old World monkeys we did not find allelic variation, but estimate one absorbance peak at medium wavelengths (527 nm), corresponding to the OPN1MW paralog, and one spectral peak at long wavelengths (560 nm), corresponding to OPN1LW paralog.

## Results and discussion

In this work, we performed phylogenetic analyses of the X-linked opsin genes in species representing all major groups of primates, with the primary goal of understanding the evolutionary origin of trichromatic color vision in apes and Old World monkeys. Additionally, we estimated the spectral sensitivity in all types of X-linked opsin sequences found in primates, using the five site rule proposed by Yokoyama and Radlwimmer^21^.

### Phylogenetic analyses suggest an independent origin of routine trichromacy in apes and Old World monkeys

*Our phylogenetic reconstruction shows that the gene tree did not deviate significantly from the expected organismal relationships. Our results showed a well-supported clade (85*.*2/0*.*9/95) containing the OPN1LW sequences of strepsirrhine primates (Fig. 1). Subsequently, the strepsirrhine clade was shown to be associated with a well-supported clade (65*.*5/0*.*9/80) containing haplorhine X-linked opsin genes (Fig. 1). Our analyses revealed the tarsier OPN1LW sequence to be related to the clade containing the anthropoid X-linked opsin genes (99*.*4/1/100; Fig. 1). The New World monkey OPN1LW clade was determined with strong support (99*.*2/1/100) to be directly related to the catarrhine X-linked opsin clade (Fig. 1)*.

**Figure 1.**
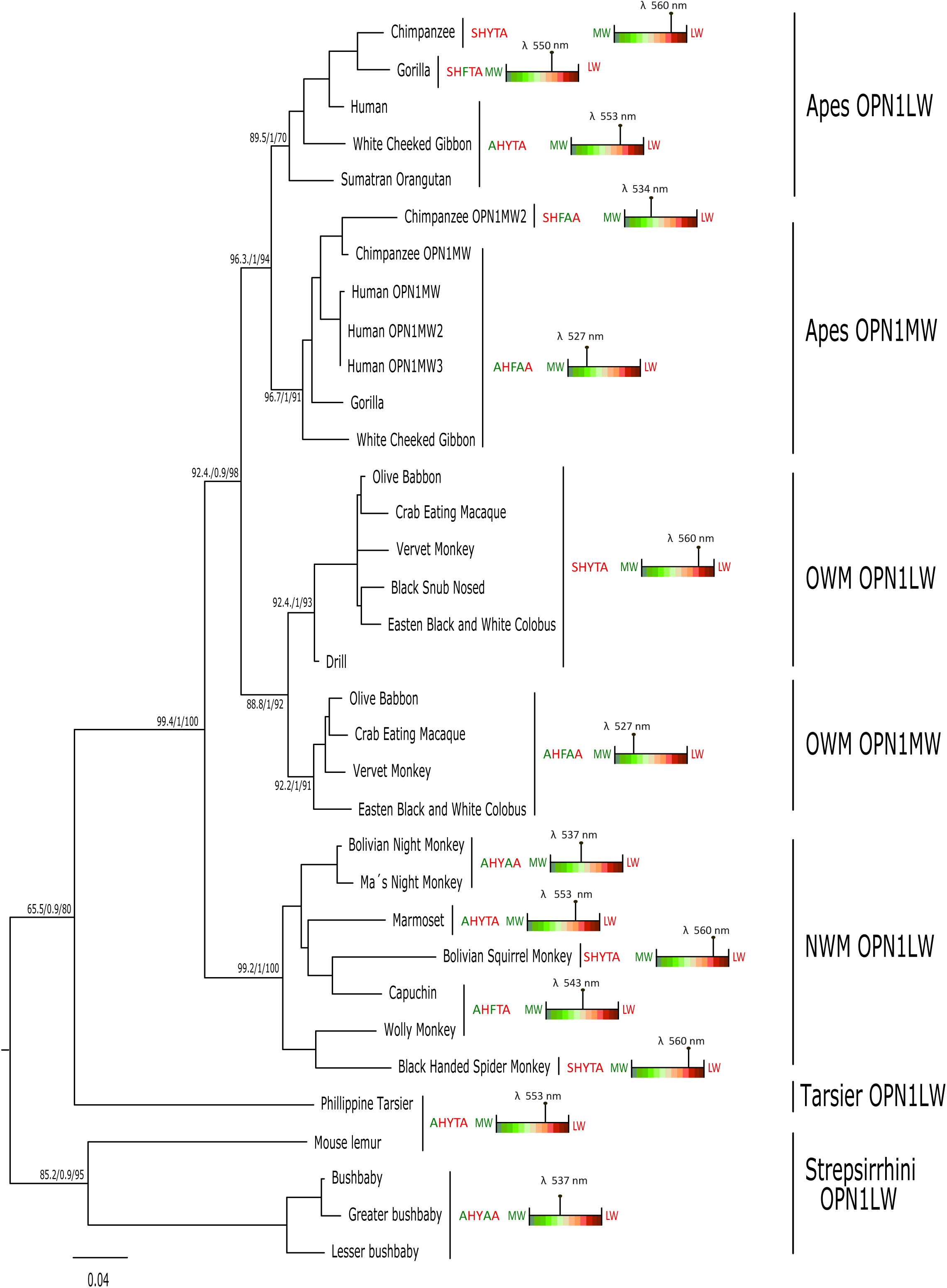
Maximum likelihood phylogram describing relationships among X-linked opsin genes of primates. Numbers on the nodes correspond to support from the Bayesian-like transformation of aLRT (aBayes test), SH-like approximate likelihood ratio test (SH-aLRT) and the ultrafast bootstrap approximation. Letters at the right hand side of the species names are the amino acid identity of the five residues (180, 197, 277, 285 and 308), which mainly predicts the maximum wavelength of absorption of the visual spectrum according to the five sites rule of Yokoyama and Radlwimmer (1999). The color rectangle represents the wavelength of the visual spectrum between the medium (green) and the long (red) range. The mark above represents the spectral pick according to the five sites rule. Cow (*Bos taurus*) OPN1LW sequence was used to root the tree (not shown).

*In contrast to other primate groups, catarrhines are characterized by having a greater diversity of X-linked opsin genes (OPN1LW and OPN1MW), an attribute that grants them routine trichromacy*^*11*^. *If the duplication event that gave rise to the OPN1MW and OPN1LW genes occurred in the catarrhine ancestor, we should expect a tree topology in which the clade containing OPN1LW sequences from apes is shown to be directly related to the clade containing OPN1LW sequences from Old World monkeys. The same pattern should be expected for the OPN1MW sequences of both primate groups. According to our results this is not the case (Fig. 1). Our gene tree suggests that routine trichromacy evolved from dichromatic color vision*^*1,22*^ *twice during the evolutionary history of catarrhines (Fig. 1). This hypothesis is supported by the fact that we discovered a clade (88*.*8/1/92) containing all paralogs found in Old World monkeys (OPN1MW and OPN1LW; Fig. 1), to be sister to a strongly supported clade (96*.*3/1/94) containing all paralogs identified in apes (OPN1MW and OPN1LW; Fig. 1). In the case of Old World monkeys, we showed a clade containing the genes located at the 5*’ *side of the cluster, i*.*e. OPN1LW, to be sister to a clade containing the genes located at the 3’ side of the cluster, i*.*e. OPN1MW (Fig. 1). This phylogenetic arrangement suggests that the duplication event that gave rise to the OPN1LW and OPN1MW genes in Old World monkeys occurred in the group’s ancestor, between 29*.*44 and 19*.*4 millions of years ago*^*23*^. *Similarly, within the apes branch we also showed a clade containing the genes located at the 5*’ *side of the cluster, i*.*e. OPN1LW, to be sister to a clade containing the genes located at the 3*’ *side of the cluster, i*.*e*., *OPN1MW. This suggests that the duplication event that gave rise to trichromatic color vision in apes occurred between 29*.*44 and 20*.*19 millions of years ago, in the group’s ancestor*^*23*^. *The independent duplications of the X-linked opsin gene in the ancestors of apes and Old World monkeys are not the only ones that occurred in the evolution of primates*^*13,18,24*^. *In New World monkeys, species of the genus Alouatta (howler monkeys) also routinely have trichromatic vision. So far there is evidence of duplicated X-linked opsin genes in at least four species (Venezuelan red howler, Alouatta seniculus; black howler, Alouatta caraya; mantled howler, Alouatta palliata; Guatemalan black howler, Alouatta pigra)*^*13,18,25*^, *suggesting that the duplication event occurred in a howler monkey ancestor. Beyond opsin genes, the independent origin of groups of genes that are associated with a specific function seems to be rather common in nature. There are several examples in which gene families (e*.*g. ß-globins, TRPV5/6 channels, GDF1/3, MHC class II genes, growth hormone) arose independently in different taxonomic groups*^*26,27,28,29,30,31,32,33*^.

*It is important to mention that although we are suggesting that duplicated copies of the X-linked opsin genes originated independently in the last common ancestors of apes and Old World monkeys, we cannot rule out alternative scenarios. A similar tree topology could be obtained if the OPN1LW and OPN1MW paralogs originated from a single duplication event in the common ancestor of apes and Old World monkeys*^*3,7,22*^. *However, we would have to assume that after duplication in each of the ancestors, these genes underwent independent gene conversion events. Furthermore, we also must assume that conversion events stopped before the radiation of both groups*.

### Variation in spectral sensitivity of primate X-linked opsin genes

Among the X-linked opsin genes, we found variation in spectral sensitivity according to the five sites rule, which predicts the maximum wavelength for absorption of the visual spectrum, based on the amino acid identity of five residues of the opsin protein^21^.

In accordance with the literature, among strepsirrhines we found two alleles (M and L) that possess different maximum wavelength of absorption (Fig. 1)^19^. The first type (AHYAA) possesses a predicted maximum wavelength of absorption at 537 nm and is present in three species: bushbaby (*Otolemur garnettii*), greater bushbaby (*Otolemur crassicaudatus*) and lesser bushbaby (*Galago senegalensis*)(Fig. 1). This estimate differs from the electrophysiological measurements (543-545 nm) performed for the greater bushbaby (*Otolemur crassicaudatus*)^15,34^. This could be explained by the fact that in addition to the 5 sites mentioned above, there are other sites that also contribute to determining the maximum wavelength of absorption, but to a lesser extent. (e.g. S116Y, I230T, A233I, Y309F)^2,5,34,35^. The second type (AHYTA) was found in the mouse lemur (*Microcebus murinus*) and has a predicted maximum wavelength of absorption at 553 nm (Fig. 1). This last estimate is similar to the expected maximum wavelength of absorption for the primate ancestor^7,19,21^. Although most strepsirrhines possess dichromatic color vision, allelic trichromacy is described but in few species (two diurnal lemurs: Coquerel’s sifaka (*Propithecus coquereli*), red ruffed lemur (*Varecia rubra*); one cathemeral lemur: blue-eyed black lemur (*Eulemur macaco flavifrons*) and one nocturnal lemur, the greater dwarf lemur (*Cheirogaleus major*))^19,36,37^. In the tarsier (*Carlito syrichta*), the identity of the five amino acid motif sequence is the same as the one found in the mouse lemur (AHYTA)(Fig. 1). This is in agreement with the results reported by Tan and Li (1999)^19^ and Melin et al. (2013)^20^ for the amino acid positions 180, 277 and 285. In the literature, it is reported that tarsier species possess an exclusive set of polymorphisms^19^. For example, the Bornean tarsier (*Tarsius bancanus*) possesses seven medium wavelength sensitive alleles (M), whereas the Philippine tarsier (*Carlito syrichta*) has 25 long wavelength sensitive alleles (L)^19^. Thus, although tarsiers have dichromatic vision, the color perception of the two species is different, similar to protanomalous and deuteranomalous dichromatic human color vision^5,20^.

On the other hand, some authors mention that it is possible the literature has underestimated the number of prosimian species with allelic trichromatic vision because many studies have been restricted to small cohorts of animals^9^. Among New World monkeys, we found four alleles, two with a maximum wavelength of absorption in the red range (560 and 553 nm), one with a medium wavelength of absorption in the green range (537 nm), and another in an intermediate range of absorbance (543 nm). In our taxonomic sampling, we did not find an allele with less spectral sensitivity described for this group (530 nm)^21,35^. The motif SHYTA with 560 nm of spectral sensitivity was found in the black-handed spider monkey (*Ateles geoffroyi*) and Bolivian squirrel monkey (*Saimiri boliviensis*), whereas the motif AHYTA with 553 nm of spectral sensitivity was found in the marmoset (*Callithrix jacchus*)(Fig. 1). We also found a green opsin phenotype with motif AHYAA in the Ma’s night monkey (*Aotus nancymaae*) and the Bolivian night monkey (*Aotus azarae*) with a spectral sensitivity peak of 537 nm (Fig. 1). In the woolly monkey (*Lagothrix lagotricha*) and capuchin monkey (*Cebus capucinus imitator*) the motif AHFTA encodes for an intermediate phenotype with a spectral sensitivity peak of 543 nm (Fig. 1). Our results are consistent with those described previously for New World monkeys, in which the identity of three amino acid positions (180, 277 and 285) are used to estimate of the maximum wavelength of absorption^14,38,39,40^, through electroretinography measurements or *in vivo* measurements of cone expression. In Old World monkeys we did not find variation (Fig. 1). The clade corresponding to the OPN1LW gene lineage possesses an amino acid motif (SHYTA) that has a maximum wavelength of absorption in the red phenotype (560nm)(Fig. 1). On the other hand, the OPN1MW gene lineage has an amino acid motif (AHFAA) that possesses a maximum wavelength of absorption in the green phenotype (527nm)(Fig. 1). This is consistent with what is reported in the literature, where it is stated that the basic genotype of the M and L opsin genes is essentially identical in Old World monkeys^41^. In the apes group, we found three alleles in the OPN1LW clade (SHYTA, AHYTA and SHFTA) with spectral ranges that vary between 560 and 553 nanometers of the visual spectrum, therefore in the red phenotype. Also, in the clade containing OPN1MW sequences, we identified two alleles (SHFAA and AHFAA) with a medium wavelength absorption (Fig. 1).

### Concluding Remarks

In the present study we show a phylogenetic analysis of the X-linked opsin genes in primates and a survey of the diversity of color vision based on the five sites rule of Yokoyama & Radlwimmer^21^. We propose that routine trichromacy, i.e. the condition of having trichromatic color vision based on having two x-linked opsin genes (OPN1MW and OPN1LW) in addition to an autosomal gene (OPN1SW), originated independently in the ancestors of apes and Old World monkeys. The selective advantage of trichromatic color vision lies in the long-range detection of either ripe fruits or young leaves against a background of mature foliage^42^. This is supported by theoretical studies of object visibility based on the colorimetric properties of natural scenes measured in forests^43,44,45^. In addition, there is recent evidence that relates the distribution and diversification of African primates with trichromatic color vision and the availability of reddish conspicuous fruits^46^, supporting the idea of the importance of trichromatic color vision in crucial aspects and coevolutionary dynamics between sensory systems and the environment. This knowledge added to the phylogenetic approach can provide important insight into the process of molecular adaptation.

## Acknowledgements

This work was supported by Fondo Nacional de Desarrollo Científico y Tecnológico (FONDECYT) grant 1160627 and Millennium Nucleus of Ion Channels Associated Diseases (MiNICAD), Iniciativa Científica Milenio, Ministry of Economy, Development and Tourism, Chile, to JCO. J.T-V acknowledges support from Fondo Nacional de Desarrollo Científico y Tecnológico (FONDECYT) postdoctoral grant 3180679.

## Material and Methods

### Sequence data and phylogenetic analyses

We used bioinformatics searches to identify and manually annotate a full complement of X-linked opsin genes in 23 primate species (Supplementary table 1). Our sampling included four strepsirrhines, mouse lemur (*Microcebus murinus*), bushbaby (*Otolemur garnettii*), greater bushbaby (*Otolemur crassicaudatus*), *and* lesser bushbaby (*Galago senegalensis*); one tarsier, Philippine tarsier (*Carlito syrichta*); seven New World monkeys, ma’s night monkey (*Aotus nancymaae*), Bolivian night monkey (*Aotus Azarai boliviensis*), Bolivian squirrel monkey (*Saimiri boliviensis*), marmoset (*Callithrix jacchus*), capuchin (*Cebus capucinus*), woolly monkey (*Lagothrix lagotricha*) and black-handed spider monkey (*Ateles geoffroyi*); six Old World monkeys, olive baboon (*Papio anubis*), easten black and white colobus (*Colobus guereza*), crab eating macaque (*Macaca fascicularis*), black snub nosed (*Rhinopithecus bieti*), drill (*Mandrillus leucophaeus*) and vervet monkey (*Chlorocebus aethiops*) and five apes, white cheeked gibbon (*Nomascus leucogenys*), sumatran orangutan (*Pongo abelii*), gorilla (*Gorilla gorilla gorilla*), chimpanzee (*Pan troglodytes*) and human (*Homo sapiens*). DNA sequences were obtained from the Ensembl database v. 98^47^ and the National Center for Biotechnology Information (NCBI) database (refseq_genomes, htgs, and wgs)^48^. Genomic pieces were extracted including the 5’and 3’ flanking genes. After extraction, we curated the existing annotation by comparing known exon sequences to genomic pieces using the program Blast2seq v.5 with default parameters^49^.

Putatively functional genes were characterized by an intact open reading frame with the canonical 6 exon/5 intron structure typical of a mammalian opsin gene. Nucleotide sequences were aligned using the L-INSI-i strategy from MAFFT v.7^50^. We used the proposed model tool of IQ-Tree v.1.6.12^51^ to select the best-fitting model of codon substitution (MGK+F3×4+R3). This approach uses a more realistic description of the evolutionary process at the protein-coding sequence level by incorporating the structure of the genetic code in the model^52^. We used the maximum likelihood method to obtain the best tree using the program IQ-Tree v1.6.12^53^ and assessed support for the nodes using three strategies: a Bayesian-like transformation of aLRT (aBayes test)^54^, SH-like approximate likelihood ratio test (SH-aLRT)^55^ and the ultrafast bootstrap approximation^56^. Cow (*Bos taurus*) OPN1LW sequence was used as an outgroup.

